# Computational Analysis of Effects of Clot Length on Acute Ischemic Stroke Recanalization under Different Cyclic Aspiration Loading Conditions

**DOI:** 10.1101/2022.04.06.487217

**Authors:** Priyanka Patki, Scott Simon, Keefe B. Manning, Francesco Costanzo

## Abstract

Acute ischemic stroke, the second leading cause of death worldwide, results from occlusion of a cerebral artery by a blood clot. Application of cyclic aspiration using an aspiration catheter is a current therapy for the removal of lodged clots. In this study, we perform finite element simulations to analyze deformation of long clots, having length to radius ratio of 2 to 10, which corresponds to clot-length of 2.85–14.25 mm, under peak-to-peak cyclic aspiration pressures of 10 to 50mmmHg, and frequencies of 0.5, 1 and 2 Hz. Our computational system comprises of a nonlinear viscoelastic solid clot, a hyperelastic artery, and a nonlinear viscoelastic cohesive zone, the latter modeling the clot–artery interface. We observe that clots having length-to-radius ratio approximately greater than two separate from the inner arterial surface somewhere between the axial and distal ends, irrespective of the cyclic aspiration loading conditions. The stress distribution within the clot shows large tensile stresses in the clot interior, indicating the possibility of simultaneous fragmentation of the clot. Thus, this study shows us the various failure mechanisms simultaneously present in the clot during cyclic aspiration. Similarly, the stress distribution within the artery implies a possibility of endothelial damage to the arterial wall near the end where the aspiration pressure is applied. This framework provides a foundation for further investigation to clot fracture and adhesion characterization.

## 1 Introduction

Acute ischemic stroke (AIS) accounts for about 87% of stroke-related deaths [13], and has been reported as a leading cause of deaths worldwide [4]. Moreover, a global increase in the aging population and accumulation of risk factors have led to an increase in lifetime risk of stroke [51]. AIS occurs when a cerebral artery gets obstructed due to a blood clot. The procedure of mechanical thrombectomy can be used for the removal of blood clots obstructing larger blood vessels, in order to increase the therapeutic window by up to 24 hours [33]. There are currently two types of mechanical thrombectomy procedures, that either use stent retrievers like Solitaire (Meditronic) and Trevo (Stryker), or aspiration catheters such as Penumbra [5, 23], which are used separately or together.

In spite of the advancement in mechanical thrombectomy devices, complete recanalization (i.e., the re-establishment of normal blood flow) is still achieved in only about 85% patients [18]. Moreover, after the procedure, there still remains a considerable chance of the patient suffering major disabilities or even death, possibly due to partial opening of the obstructed blood vessel, or due to the large time required for reperfusion after the procedure ([15, 18]). One of the strategies implemented in order to maximise the number of positive outcomes is the implementation of cyclic suction pressure variation while using aspiration catheters [43, 44]. A general observation is that the use of cyclic aspiration as opposed to static (application of constant suction pressure) reduced the time for removal of clots and also increased the overall clearance [43].

Numerical modelling can give preclinical assessment of medical devices and thus, can aid in the understanding of the procedure limitations and further aid in the development of new procedures and devices. In this regard, numerical investigation of aspiration thrombectomy has been done via computational fluid dynamics (cfd), lumped parameter model (lpm) and finite element method (fem) methods.

Shi *at al*. [42] performed a cfd analysis of blood flow around the catheter tip during aspiration. They evaluated the relationship between aspiration pressure and clot-catheter distance during aspiration focusing exclusively on the flow around the catheter tip neglecting to model the actual clot. Pennati *et al*. [35] and Soleimani *et al*. [45] performed a two-phase fluid cfd analysis to study the factors affecting the suction of clots as a function of catheter design during aspiration of clots in coronary arteries. Blood flow in the Circle of Willis during aspiration was studied using a coupled 1D–3D volume of fluid technique [31]. These cfd studies assume the clot to be fluid-like and do not include any modeling of the clot-artery interface.

The Romero and Talayero groups focus on the use of Bond Graph based lpms for studying aspiration [38–40, 46]. In a recent study, they expanded use of this technique for the calibration of a coupled fem–cfd approach [41, 47]. They used surface tension as the only interfacial force to model the clot-artery surface adhesion, without providing any experimental validation.

A fem study evaluating the performance of two aspiration catheters involved the modeling blood clots as a combination of a solid fibrin network consisting of randomly distributed branch points embedded in the blood plasma (a fluid) [7]. They modeled the effect of aspiration pressures on the blood clots by tracking the plasma velocity and fibrin deformation, while holding the position of the branch points near the arterial wall fixed. Hence, they did not incorporate any clot–artery interfacial interactions as the clots were assumed to be sticking to the arterial wall at all times. In a more recent fem study, a new constitutive clot model that accounts for plastic deformation of clots was developed and used for the analysis of aspiration thrombectomy [12]. The authors consider aspiration pressure applied to a free-standing clot through a catheter and do not consider clots occluding arteries in their computational model.

Recanalization rates are highly dependent on the mechanical structure of the clot as well as the contact mechanics at the interface of the clot and inner wall of the occluded artery [53]. Hence, in order to accurately model the clot-artery interface, our group formulated a computational framework with an initial validation using benchtop experiments [14]. The framework makes use of accurate models to describe the behavior of clots and arteries and the clot-artery interface. In order to model the interface, we use the concept of cohesive zone (cz) modelling (c.f. [8]), where the interface is modelled as a bio-film having negligible inertia. In previous studies, we used the developed computational framework to analyze the use of cyclic aspiration for the removal of clots from cerebral arteries, while treating the artery as rigid [14] and deformable [34].

In the current work, we expand our analysis of cyclic aspiration by studying its effectiveness and limitations in the removal of long clots. Clinical literature indicates poor clinical and safety outcomes of aspiration of long clots [53, 54]. In order to increase our chances of successful first-pass retrieval of long clots, it is important to understand the underlying failure mechanisms at play during cyclic aspiration of long clots. The current study is, to the best of our knowledge, the first such numerical investigation and has important implications in terms of the various failure mechanisms in the clot during cyclic aspiration and also points towards the possible post-surgery damage to the occluded artery.

## 2 Numerical framework

We adopt the computational framework used in [34] to simulate a clot lodged in an hyperelastic artery. The initial geometry of the system is shown in Fig. 1(a). The clot is assumed to be a solid right circular cylinder, and the artery is assumed to be a concentric, hollow right circular cylinder. The clot is initially in contact with the artery along the lateral surface and is exposed to blood at the bottom surface. A negative pressure is applied at the lower end of the clot through an aspiration catheter. The end where the aspiration pressure is applied will be henceforth referred to as the proximal end, and the opposite end will be called the distal end. In reality, the catheter is a long, hollow tube with finite wall thickness. Hence, for modeling purposes, we assume that the diameter over which the suction pressure is applied is smaller than the diameter of the clot itself. We use cz theory to simulate the interface between the clot and artery. Our computational system thus, consists of three components: a hyperelastic artery, a nonlinear viscoelastic solid clot, and a cohesive zone (cz) that models the clot-artery interface.

**Figure 1:**
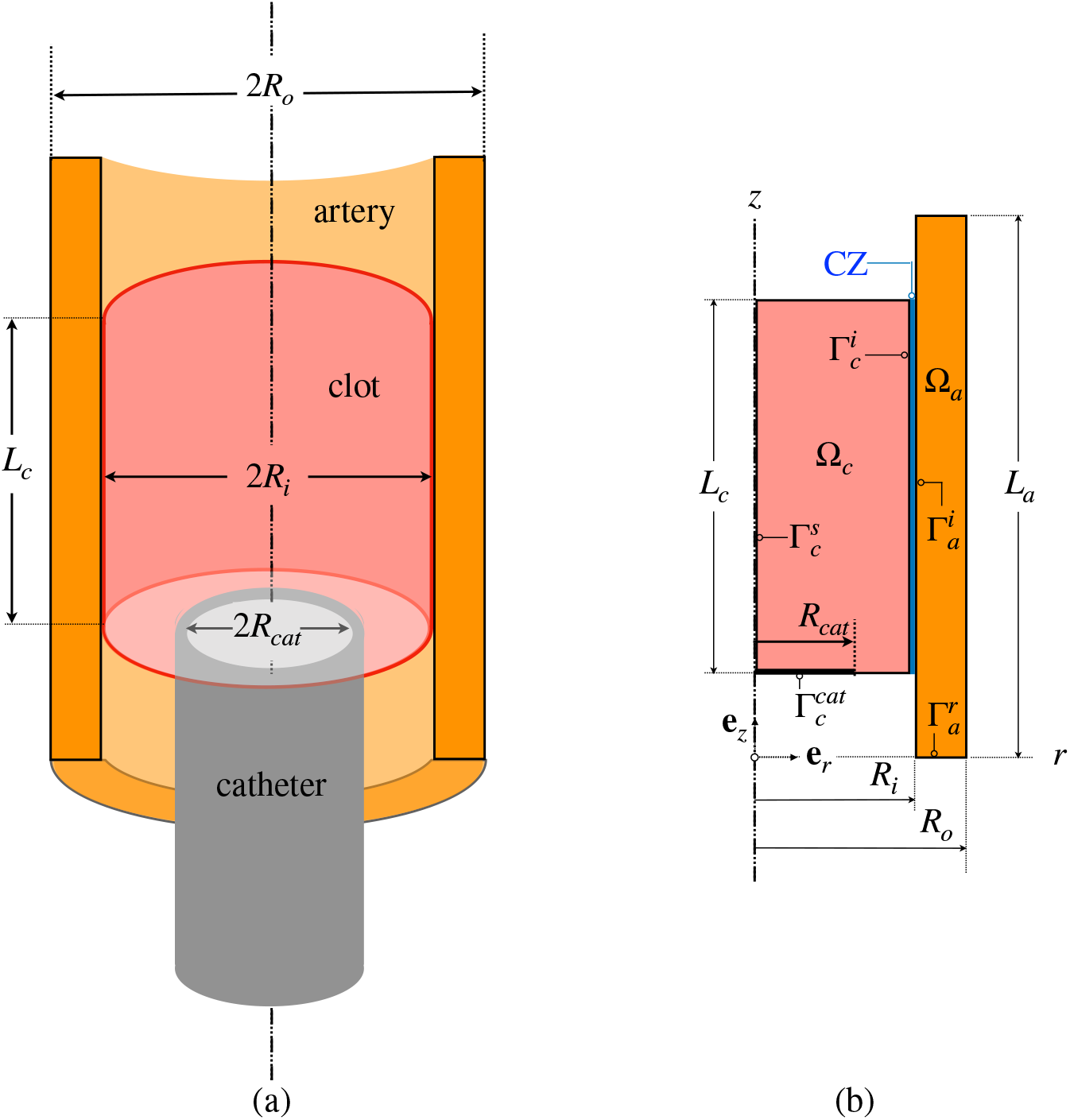
(a) Initial geometry of the system comprising of a right cylindrical clot, concentric with an artery. Aspiration pressure is applied through a catheter having inner diameter 2*R_cat_* at the lower end of the clot. (b) Reference (initial) configuration showing the 2D axisymmetric computational domains. Unit vectors along the radial and axial direction are denoted by **e**_*r*_ and **e**_*z*_ respectively. For subscripts *a* and *c* denoting the artery and clot, respectively, the reference domains are denoted by Ω_*c*_ and Ω_*a*_; The interface boundaries, which are in direct contact with the cz, are 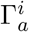 and 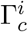 Motion is restricted to radial-only is applied at the arterial boundary 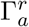. Axial symmetry boundary conditions are applied at the clot boundary 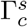, and a pressure boundary condition is applied on the boundary 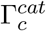, which is a subset of the proximal clot boundary, and has length *R_cat_*.

### 2.1 Notation and Kinematics

For each component in our system, we define two configurations, the reference configuration and the deformed (or current) configuration. For our simulations, we assume that the reference configurations of the clot and artery coincide with their initial configurations. We label the particles in the reference and current configurations by **X**_*i*_ and **x**_*i*_, respectively. The corresponding gradient and divergence operators are given by ∇_**X**_*i*__, ∇_**x**_*i*__, ∇_**X**_*i*__ · and ∇_**x**_*i*__ ·, respectively. The subscript *i* can take the values *a* or *c* representing the artery and clot, respectively. Additionally, quantities pertaining to the cz carry the subscript *cz*. The reference and current configurations are mapped by the diffeomorphism given by **x**_*i*_ = χ_*i*_(**X**_*i*_, *t*), where *t* is time. We define the displacement, deformation gradient for the mapping, its Jacobian, and the left and right Cauchy-Green tensors by **w**_*i*_ = **x**_*i*_ – **X**_*i*_, F_*i*_ = ∇_**X**_*i*__**x**_*i*_, *J_i_* = determinant(F_*i*_), B = FF^T^ and C = F^T^F, respectively. The material time derivative of any quantity *α* is denoted by 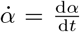. The material velocity for component *i* is defined as 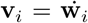. The gradient of velocity and its symmetric part are denoted by L_*i*_ = ∇_**x**_*i*__**v**_*i*_ and 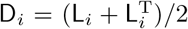. The tensor product of two vectors, **r** and **n** is denoted by **r** ⊗ **n**.

Referential unit vectors along the radial and axial directions are denoted by **e**_*r*_ and **e**_*z*_. We define by ***δ***, the cz opening displacement, given by ***δ*** = **w***_a_* – **w**_*c*_ + *θ**e**_r_*, where *θ* is a small nominal value for the cz thickness. *δ* and **e***_cz_* denote the magnitude of ***δ*** and unit vector along ***δ***, respectively. For our simulations, we assume that the initial and undeformed configurations coincide with the referential configurations for the clot and artery. The initial configurations for our system are shown in Fig. 1(b), where Ω_*a*_ and Ω_*c*_ denote the computational domain for the artery and clot, respectively.

### 2.2 Constitutive models

For this study, we largely follow the constitutive models used in [34], which we describe in short here.

#### Artery

we model the artery as an incompressible, hyperelastic solid [21]. The Cauchy stress for the artery is given by,

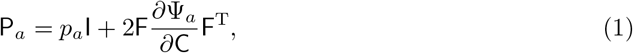

where *p_a_* is the Lagrange multiplier for the enforcement of the incompressibility constraint, and Ψ_*a*_ is the strain energy per unit referential volume of the artery. We consider elastic contributions of the media and fibers. We then define Ψ_*a*_ as follows [9],

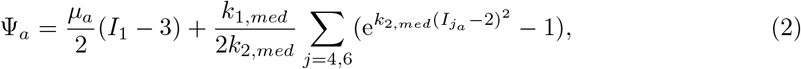

where *μ_a_, k*_1,*med*_, and *k*_2,*med*_ are material constants. The fibers within the artery are assumed to be oriented along two directions. The unit vectors along these fiber orientations are denoted by **m**_1_ and **m**_2_ that make angles *θ*_*fib*_1__ and *θ*_*fib*_2__ with the radial direction. Then, for A_1_ = **m**_1_ ⊗ **m**_1_ and A_2_ = **m**_2_ ⊗ **m**_2_, we have *I*_1*a*_ = trace(C_*a*_), *I*_4*a*_ = trace(C_*a*_A_1_), and *I*_6*a*_ = trace(C_*a*_A_2_).

#### Clot

We model the clot as an incompressible Kelvin-Voigt solid (c.f. [50]), which is representative of mature clots. The Cauchy stress for the clot is given by

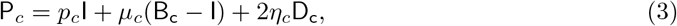

where *p_c_* is a Lagrange multiplier for incompressibility, *μ_c_* and *η_c_* are constant shear and elastic moduli for the clot, and I is the 2^nd^ order identity tensor.

#### Cohesive zone

Since the interface between the clot and the inner arterial wall can be expected to have behavior similar to that of the clot itself, we adopt the cz model used in [14] and [34], wherein the cz is modeled as a nonlinear viscoelastic fluid. The cz is modeled as a surface having negligible inertia, which imparts an equal and opposite traction distribution on the clot and inner arterial wall. We assume that the traction exerted by the cz on the clot surface is given by,

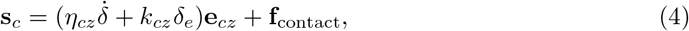

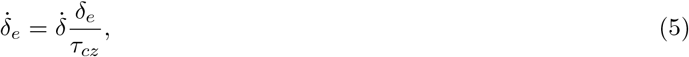

where *τ_cz_* is a characteristic relaxation time constant and *k_cz_* and *η_cz_* are material constants. **f**_contact_ is an elastic contact force, which acts perpendicular to the contact surfaces only when the clot presses against the artery and is defined as,

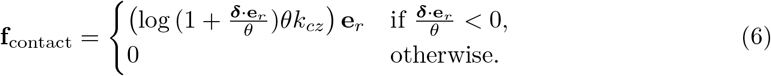

For constant values of *k_cz_* and *η_cz_*, the cz model given by Eqs. (4)–(5) allows unrestricted opening of the interface under progressively increasing load. If we assume the cohesive interface to be made up of microscopic fibrils of length *l*, then the damage accumulation can be thought to be triggered by the failure of the fibrils (c.f. [14, 34]). When *δ* exceeds a critical value, *δ_cr_*, damage accumulation can be thought to be triggered. Hence, for this study, we consider two cz models — one with constant constitutive properties of the cz and another which accounts for the progressive damage accumulation of the interface due to the pullout of fibrin. This gives us the following modified model for the constitutive parameters,

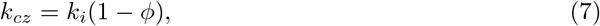

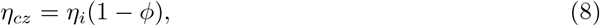

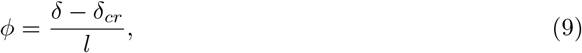

where *k_i_* and *η_i_* are the undamaged values of *k_cz_* and *η_cz_*, respectively and *ϕ* is a damage variable.

### 2.3 Initial-boundary value problem definition

The computational domain for the problem is shown if Fig. 1.

#### Mass and momentum balance equations

The balance of momentum and mass (incompressibility constraint) equations for the artery and clot, in the referential configurations, Ω_*i*_, are given by,

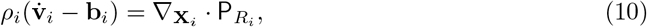

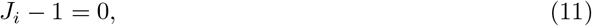

for *i* = *a, c*, where *ρ_i_* and **b**_*i*_ are the density and body force per unit referential volume, respectively, and P*_R_i__* is the first Piola-Kirchhoff stress, defined as 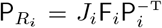, for component *i*, respectively. Due to incompressibility, the referential and current densities are identical.

#### Boundary and initial conditions

Referring to Fig. 1(b), the following boundary and initial conditions apply:

- ***For the clot*:**

– traction boundary condition given by Eq. (4) at boundary 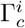.
– effect of catheter: for our simulations, we assume that the clot is not in direct contact with the catheter surface. Hence we only model the applied aspiration pressure (traction) boundary condition at the boundary, 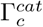, where the applied pressure, *p_asp_*, as a function of time is given by,

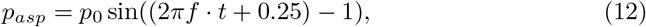

where *f* is the frequency of aspiration cycles and *p*_0_ is the peak-to-peak aspiration pressure.
– axisymmetry boundary conditions at the boundary 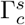,
- ***For the artery*:**

– traction boundary condition given by Eq. (4), where **s**_*a*_ = –**s**_*c*_ at the boundary 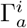,
– restricted: axial-only motion for the boundary 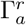, given by,

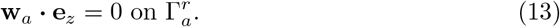
- ***Initial conditions*:**

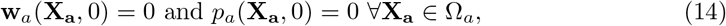

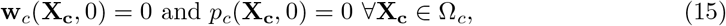

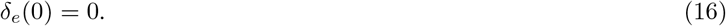

#### Parameter values

Referring to Fig. 1(b), we chose clot length, *L_c_*, as 2*R_i_* = 2.85 mm, 5*R_i_* = 7.125 mm, 7*R*_*i*_ ≈ 10 mm, 10*R*_*i*_ = 14.25 mm for different cases. For our computations, the length of the artery was set to *L_a_* = *L_c_* + (*R_o_* – *R_i_*) and was placed radially symmetric with respect to the clot. The peak-to-peak aspiration pressures were chosen as 10, 20, 30 and 50 mmHg for different cases. All the other values of constitutive and geometric parameters are given in Table 1. We refer readers to [34] and the references therein for a discussion on the choice of parameter values.

**Table 1:**
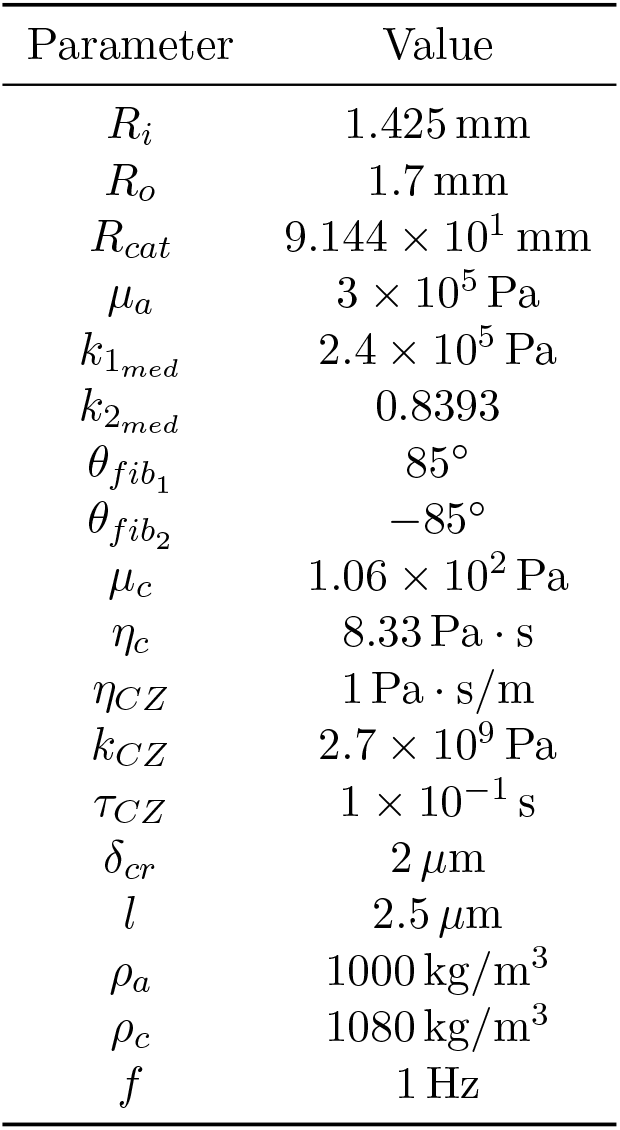
Geometric and constitutive parameter values

### 2.4 Finite element implementation

For *i* = *a, c*, we solved the initial boundary value problem (ibvp) given by Eqs. (10), (11) over the domains Ω_*i*_, along with Eq. (5), for the parameters and constitutive relations, and initial and boundary conditions discussed previously. We used a Lagrangian framework for both, the artery and the clot. The weak formulation corresponding to the ibvp was implemented in an fem framework and a modified version of the numerical stabilization scheme in [27] for the hyperelastic artery. The clot formulation was based on the formulation for complex fluids given in [22].

The resultant numerical model was implemented in COMSOL Multiphysics®. We used the ‘Weak form PDE’ interface within the Mathematics module for the clot-artery-cohesive zone formulation. Lagrange linear and discontinuous Lagrange elements were used for displacements and incompressibility multipliers, respectively. We discretized the solution domain using uniform quadrilateral elements of size 1.718 × 10^−2^ mm, based on our previous mesh refinement study, [34]. The initial value problem corresponding to the evolution of *δ_e_* was solved using the Boundary ODE interface. Second order Gauss point data shape functions were used for *δ_e_*.

Second order, variable time step backward differentiation formulas (BDF) [36], with a maximum time step size of 0.001 s was used for implicit time stepping. The resultant system of equations was solved using the non-linear solver IDAS [20]. We used a two-step segregated solver, and MUMPS [1, 2] was chosen as the linear solver for each segregated step. The first segregated step was for the evaluation of the stabilization parameter for which cubic bubble shape functions were used. The second step was for the evaluation of the solution variables, **w**_*a*_, **w**_*c*_, *p_a_, p_c_* and *δ_e_*. For the damage accumulation model, an additional segregated step for the point-wise determination of maximum *δ* was implemented.

## 3 Results

### 3.1 Deformation of clot and artery

For the computational problem discussed in the previous section, we first analyze the deformation profile of the clot and the artery. The results presented in Figs. 2 and 3 show the deformation profiles using the cz model without and with damage accumulation, respectively, for a peak-to-peak aspiration pressure of 10 mmHg. In the figures, positive radial displacement implies displacement away from the center-line and is indicated by red color on the color bar. As we progressively move towards blue color on the color bar, the radial displacement continues to decrease. Blue color indicates negative radial displacement i.e., displacement towards the center-line.

**Figure 2:**
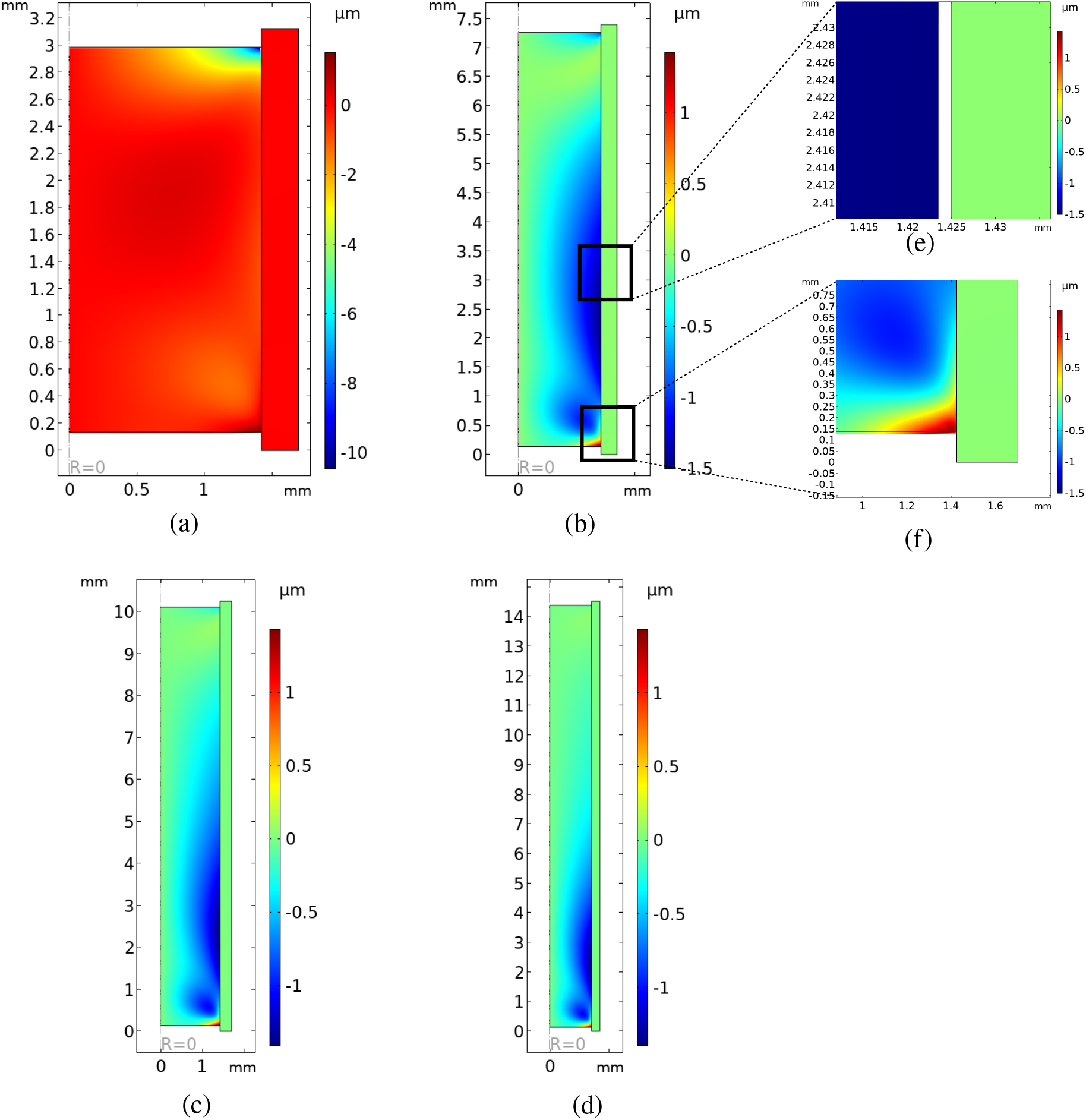
Deformation profiles using cz model without damage accumulation. Color bars indicate radial displacements, where positive and negative implies away from and towards the symmetry line, respectively. The lengths of the clots are *L_c_* = (a) 2*R_i_*, (b) 5*R_i_*, (c) 7*R_i_* and (d) 10*R_i_*, respectively, for a peak-to-peak pressure of 10 mmHg. The results are presented at time = 10 s, at the end of 10 complete aspiration cycles. The inset figures show (e) a zoomed-in view near the clot-artery interface, where the separation of the clot from the artery can be clearly observed, and (f) the location near the distal end where the clot impinges on the artery, which is marked by positive radial displacement (red color on the color bar), for a representative clot length of *L_c_* = 5*R_i_*.

**Figure 3:**
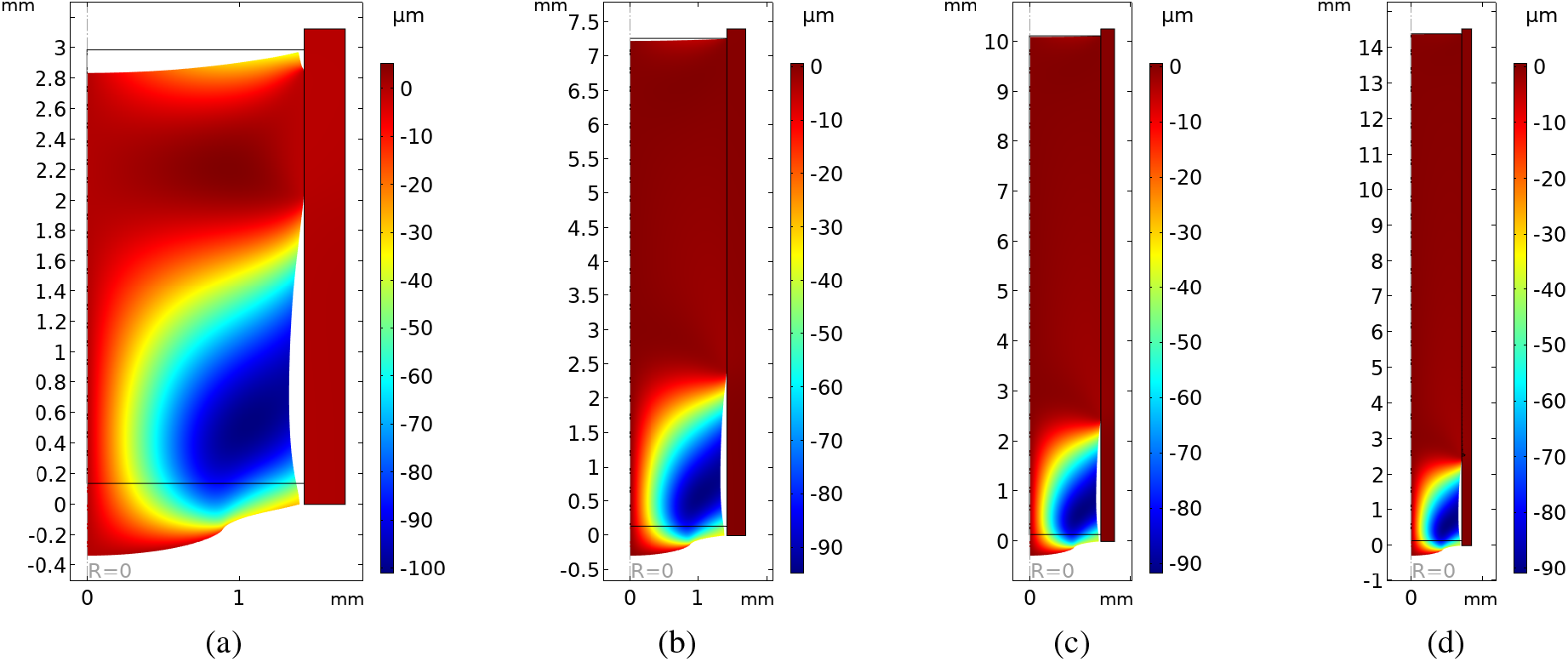
Deformation profiles using cohesive zone model that accounts for damage accumulation. Color bars indicate radial displacements. The lengths of the clots are *L_c_* = (a) 2*R_i_*, (b) 5*R_i_*, (c) 7*R_i_* and (d) 10*R_i_* respectively, for a peak-to-peak pressure of 10 mmHg. The black border shows the initial configuration of the system. The results are presented after complete failure of the cohesive interface.

As indicated by the values of *μ_a_* and *μ_c_*, the artery has higher stiffness as compared to the clot. In accordance with this fact, we observe that the artery experiences negligible deformation as compared to the clot under the action of cyclic aspiration. Aspiration of shorter clots leads to separation of the clot from the artery at the distal end [34]. However, in the case of longer clots, we observe that the clot separates from the artery somewhere between the proximal and distal ends. The absolute axial location of separation remains approximately the same (around 3 mm without damage accumulation and around 1 mm for cz model accounting for damage accumulation) for all the cases. Moreover, the point of separation is much closer to the proximal end (around 1 mm) for the cz model accounting for damage accumulation, as compared to the the case when using the cz model without damage accumulation (separation around 3 mm). As was shown in the case of short clots, we observe localized impingement of the clot against the arterial wall at the proximal end for longer clots as well, as seen in Fig. 2(f). Since the nature of deformation profiles are similar for both models and for various clot lengths, without loss of generality, we will be discussing results pertaining to one indicative case, for *L* = 5*R_i_*, using the cz model that ignores damage accumulation only, for the rest of our discussion.

### 3.2 Distribution of principal stresses

Fig. 4 shows the distribution of maximum principal stresses within the artery and clot. As observed in Figs. 4(b) and (c), there is a high concentration of principal stresses within the artery at the proximal end, where the clot presses against the inner wall of the artery.

**Figure 4:**
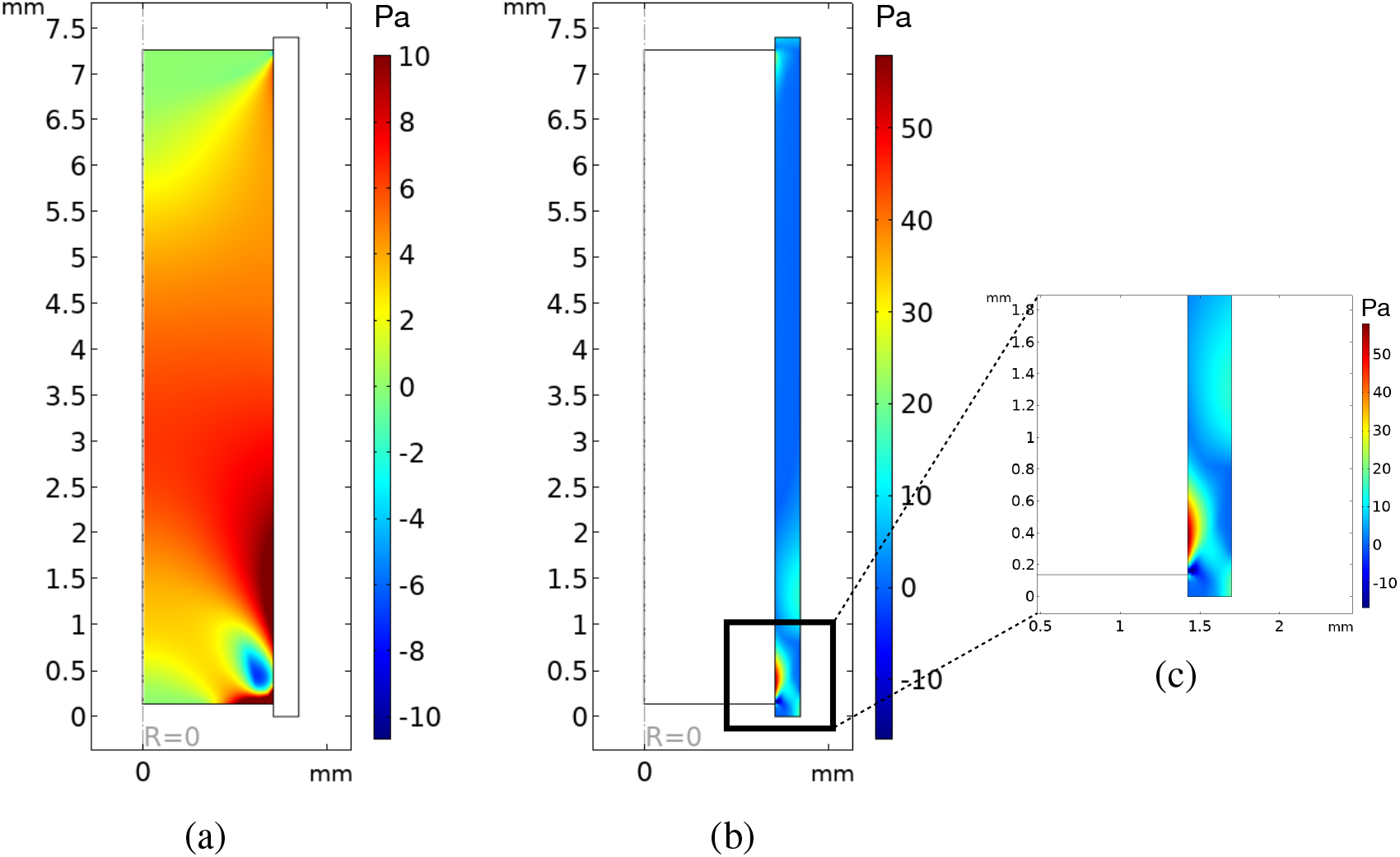
Maximum principal stress distribution: (a) for clot, (b) for artery and inset (c) shows a close-up of the principal stress distribution in the artery near the proximal end. All the results are shown for the case of *L* = 5*R_i_*. The color bar indicates stress value in Pa.

The maximum principal stress distribution observed in the clot in Fig. 4(a) is consistent with the displacement plots (Fig. 2) and indicates a possible tensile mode of clot failure.

### 3.3 Effect of pressure variation

Next, we analyzed whether changing the loading conditions in terms of change in aspiration pressure can have an effect on the separation location of the clot. The deformed configurations, along with radial displacement distributions for peak-to-peak aspiration pressures of 20, 30, and 50 mmHg are shown in Fig. 5(a), (b), and (c), respectively. As seen in Fig. 5(a)–(c), we observe that, keeping all the parameters the same, if we increase the peak-to-peak aspiration pressure then the axial location of the point where maximum separation of the clot from the artery occurs remains almost the same. Moreover, with an increase in peak pressure, we observe a deeper impingement of the clot against the arterial wall at the proximal end.

**Figure 5:**
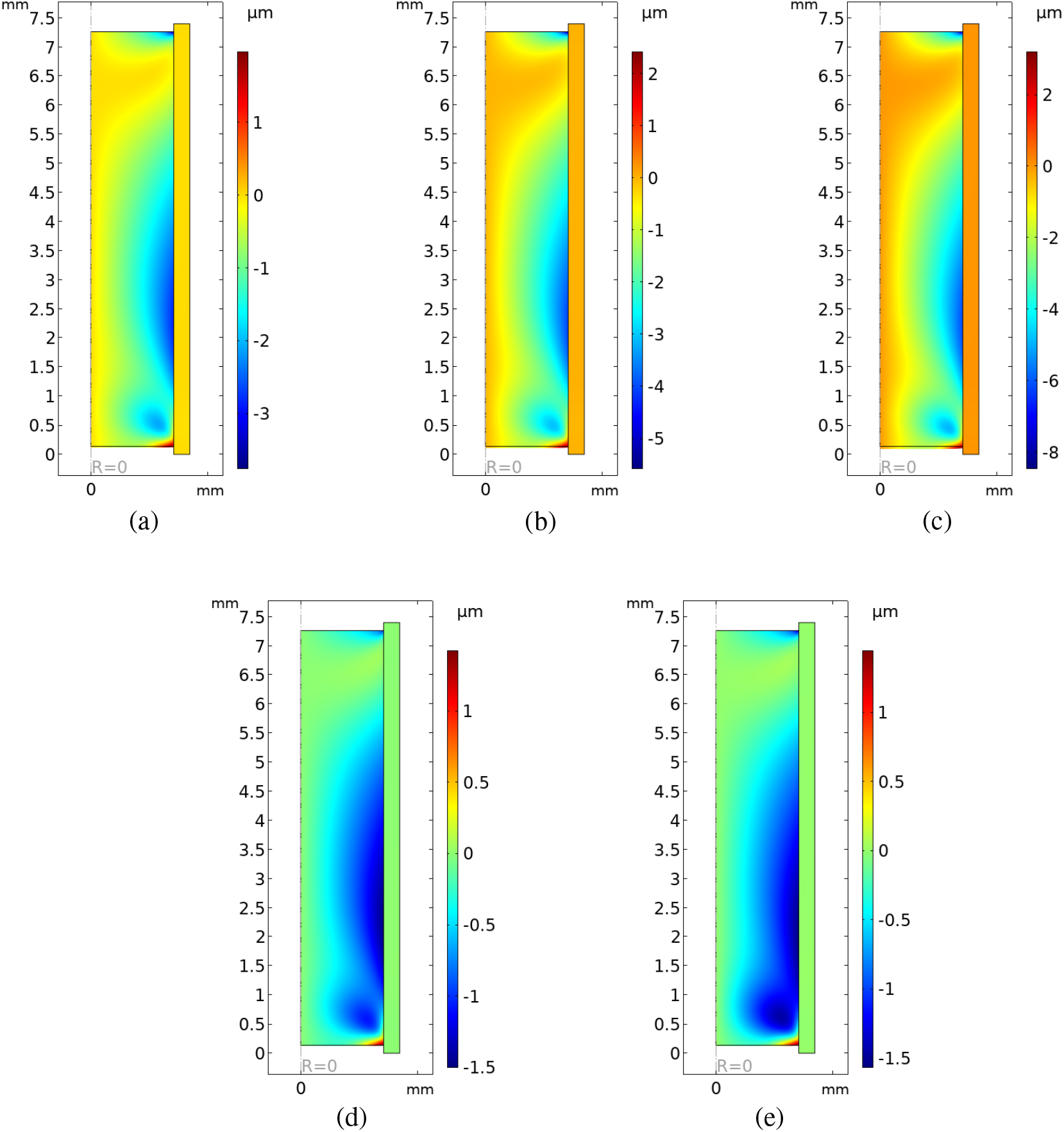
Effect of peak-to-peak pressure variation: for *L* = 5*R_i_*, for peak-to-peak pressures of (a)20, (b)30, and (c)50mmHg, for a cyclic aspiration frequency = 1 Hz. For a peak-to-peak aspiration pressure of 10 mmHg, deformation profile in the case of aspiration frequency of (d)0.5 Hz and (e)2 Hz. The color bar indicates radial displacement.

Finally, we also studied the effect of changing aspiration frequency on the deformation profile. As observed in Fig. 5(d) and (e), we observe that changing cyclic aspiration frequency has no significant effect on the location of detachment of the clot from the artery.

## 4 Discussion

Recent clinical literature shows poor performance of aspiration thrombectomy in the first passretrieval of long clots [54], possibly due to the increased clot-artery surface interactions in long clots [53]. In order to better understand the limitations posed by aspiration thrombectomy, and to develop more effective first pass strategies for the retrieval of long clots, it is essential to study the underlying mechanics of the clot and the clot-artery surface interactions during the procedure. In this regard, very few groups have developed numerical models for aspiration thrombectomy [7, 12, 31, 35, 38–42, 45–47]. However, none of these studies focus on the dependence of aspiration outcomes on clot-length, and do not include any modeling of the clot-artery interface, which is essential for understanding the limitations of the procedure. In this study, we numerically investigate the deformation and failure mechanisms of long clots during cyclic aspiration by modeling the clot as a viscoelastic solid clot, the artery as a hypereleastic solid, and the clot-artery interface as a nonlinear viscoelastic cohesive zone, following the framework in [14] and [34].

An important observation from our results is that while shorter clot separation from artery starts and propagates from the distal end [34], clots having length significantly greater than radius separate somewhere between the proximal and distal ends, as observed from Figs. 2 and 3. While the interfacial failure is initiated at an intermediate location between the proximal and distal ends, simultaneously, the clot itself exhibits a ‘conical’ distribution of maximum principal stresses, as observed in Fig. 4. Such a high concentration of maximum principal stresses indicates that eventually, the clot might undergo tensile failure in this zone. This failure cannot be observed directly in the current simulations, as our numerical framework currently does not include any model for the fragmentation of the clot within itself. However, our results show that various modes of failure might occur concurrently in the clot-artery system under cyclic aspiration.

Greater clot length directly translates into an increased clot-artery contact surface area leading to greater frictional and adhesive forces which in turn, determine how ‘sticky’ the clot is [53]. Our cz model for the clot-artery interface interactions incorporates these interaction forces between the contact surfaces. The fact that the detachment of the clot from the artery does not start from the distal end, coupled with the high principal stress distribution within the clot indicates the possibility that the distal part of the clot will remain stuck to the artery while a part of it will break apart and get removed by the aspiration catheter. In clinical scenarios, this might lead to incomplete recanalization of vessels obstructed with long clots, with a possibility that the fragments of the clot might travel with the blood flow that might lead to distal embolization i.e. the occlusion of another artery by the traveling clot. These inferences are in agreement with the recent THERAPY* trial that shows worse clinical and safety outcomes for intra-arterial aspiration thrombectomy for arteries occluded with long clots [54].

We also observe from the displacement plots that local impingement of the clot against the inner arterial wall (arterial endothelium) occurs at the proximal end, as was also observed for shorter clots [34]. Since the clot presses against the artery, this translates into a high concentration of maximum principal stresses occurring in the artery at the point of impingement, as observed from Fig. 4. Since the aspiration surgeries are carried out under unanaesthetized conditions, this might indicate that the patient might experience pain at the point of impingement during the surgery. The impingement of the clot would also indicate that, as the clot gets eventually removed, the arterial endothelium might get dragged with the clot, possibly resulting in damage to the artery. This kind of deviced-induced endothelial damage was previously reported in an *in vitro* study for aspiration as well as stent retrievers [48], and was also previously predicted in a numerical study involving two Penumbra aspiration catheters [7].

Procedural manipulations can lead to complications such as compaction of clots which in turn affect clot-artery interactions which can make clot retrieval in the next pass harder [17, 53]. Similarly, multiple passes might lead to a risk of clot fragmentation and subsequent distal embolization (occlusion of an artery due to the migration of the fragment of the original clot) [11, 17, 23, 25]. Hence, we investigate weather changing aspiration loading conditions, such as peak-to-peak pressure or cyclic aspiration frequency could help in the successful first-pass recanalization for lodged long clots. However, as observed in Fig. 5(a)–(c), increasing peak-to-peak pressure might not be the best option to obtain complete recanalization for lodged longer clots. In fact, higher impingement of the clot would imply higher stresses in the artery, which in turn might lead to higher pain and endothelial damage during the procedure. Similarly, changing the cyclic aspiration frequency had no significant effect on the location of detachment of the clot from the artery, as observed from Fig. 5(d) and (e).

Our study, in concurrence with the THERAPY clinical trial [54] indicates that direct aspiration alone might not be a sufficient single-pass clot retrieval procedure for arteries occluded with long clots. In this context, while some clinical studies indicate the ‘pinning technique’, involving a combined use of stent retriever and local aspiration to increase retrieval force on clots [10, 28], another study shows that aspiration-assisted stent retrieval is comparable within a similar outcome range with direct aspiration in terms of safety and efficacy [32]. Our study also sheds light on the underlying cause behind this, such as possible fracture of long clots due to tensile failure of the clot. Thus, altering the mechanical structure or properties of the clot through pharmacological interventions in combination with mechanical thrombectomy might lead to favourable outcomes by first softening the clot and then removing it (c.f., e.g., [16, 49, 53, 55]). While such combinations seem promising, data in this area are limited, and more targeted clinical studies involving specific combinations of thrombectomy procedures with specific pharmacology are required for efficient first-pass retrieval of long clots.

There is a significant lack of data on the mechanical properties of arterial tissues, especially for the separate layers of vessel walls, due to several practical reasons [37]. Moreover, the data on cerebral arteries is even limited, due to a lack of pragmatic mechanical catheterization [26]. In this context, several previous numerical studies on aspiration have modeled the arterial wall as rigid [7, 14, 35, 42]. The Romero and Talayero groups have used a simplified, linear elastic model for the artery [38–41, 46, 47]. For the current study, we have used the nonlinear hyperelastic model in [21], which was also used in our previous study, [34]. In the literature, we find large variation with respect to the thickness of the arterial adventitia layer. For example, certain studies claim an adventitia thickness of the order of several micrometers (c.f. [29, 30]), whereas certain studies claim thickness of the same order of magnitude as the media (c.f. [52]). From a computational modeling perspective, a very thin adventitia layer would require it to be treated as a membrane, as opposed to a layer with finite thickness. Under these considerations, for the present study, we have neglected the contribution of the adventitia. With the contribution of the intima and media itself, we observe that the artery is several orders of magnitude stiffer as compared to the clot. Hence, the results presented in this study are conservative estimates of the clot-artery response to applied cyclic aspiration.

We use a Kelvin-Voigt type solid to model the clot, which is representative of more mature clots. The constitutive response of a clot depends on a number of factors such as its age and platelet contraction. For example, coarsely ligated clots tend to have a more fluid-like behavior as described in [3]. Similarly, platelet contraction is known to significantly affect clot microstructure and mechanical properties such as stiffness [24]. Clot-artery interfacial friction is also known to be dependent on the fibrin content in the clots [19, 53] and is shown to affect recanalization rates [6, 53]. Thus, the mechanical behavior of the clot and its effects on recanalization rates during mechanical thrombectomy in itself constitutes a separate study, and would be undertaken in the future. Additionally, we also wish to incorporate models to describe fragmentation behavior of the clots. Similarly, we would like to model the contact mechanics between the catheter and the clot when the two are in contact with each other. The current study serves as a foundation to which we plan to add higher complexities in terms of clot fracture and adhesion characterization to better understand aspiration and other thrombectomy procedures for AIS.

## 5 Conclusions

In this study, we present computational results pertaining to removal of lodged clots through the application of cyclic aspiration. Unlike smaller clots, we observed that in the case of clots having length significantly larger than the radius (5 times the radius or more), the clots detach from the artery somewhere between the proximal and distal ends. Although we have not incorporated a clot fragmentation mechanism, this indicates that application of cyclic aspiration might lead to the breaking of larger clots. Clinical implications of our study are that in order to completely recanalize arteries lodged with larger clots, either multiple cyclic aspiration passes, or a use of a combination of different mechanical thrombectomy procedures might be required. Our study also sheds some light on the possible failure mechanisms in the clot-artery system under cyclic aspiration. The distribution of principal stresses indicates interfacial failure of the clot-artery interface and the associated possibility of damage of the arterial endothelium. Moreover, we observe that long, mature clots might undergo tensile failure leading to their fragmentation under cyclic aspiration.

## 6 Acknowledgements

This work was funded by NIH grant HL146921.

## 7 Conflicts of Interest

The authors declare they have no conflicts of interest.

## Footnotes

* THERAPY (The Randomized, Concurrent Controlled Trial to Assess the Penumbra System’s Safety and Effectiveness in the Treatment of Acute Stroke; NCT01429350) was a randomized trial of a combination of aspiration and IV r-tPA versus IV r-tPA alone in large-vessel stroke patients with thrombus length measurement ≥ 8 mm.

## Notes

### Competing Interest Statement

The authors have declared no competing interest.

